# The resolvin D and E biogenesis pathway regulates senescence and ageing

**DOI:** 10.64898/2026.01.26.701588

**Authors:** Mario Mira-Carnicer, Marta Menéndez-García, Antonio Merino-Navarro, Celia Palomino-Lozano, Cristina Antón-Barros, Ignacio Palmero, Andrea Malaspina, Jorge Montesinos, Ana O’Loghlen

**Affiliations:** Epigenetics & Cellular Senescence Group, Biomedicine Department, Biological Research Centre Margarita Salas (CIB), Spanish National Research Council (CSIC), Calle Ramiro de Maeztu 9, 28040 Madrid, Spain; Lipid Regulation of Immunometabolism Lab, Biomedicine Department, Biological Research Centre Margarita Salas (CIB), Spanish National Research Council (CSIC), Calle Ramiro de Maeztu 9, 28040 Madrid, Spain; Instituto de Investigaciones Biomédicas “Sols-Morreale” CSIC-UAM, Madrid, Spain; Queen Square Motor Neuron Disease Centre, Neuromuscular Department, Institute of Neurology, University College London, London, United Kingdom

## Abstract

Ageing is considered as a process were molecular, cellular and tissular function is impaired. One classic cellular phenotype that increases during ageing is cellular senescence. Upon senescence, the cells stop proliferating and release a variety of cytokines, chemokines and extracellular vesicles. However, the implication of biomolecules derived from lipids such as resolvins are not well characterised in senescence and ageing. Here, we find that the resolvin E and D biosynthesis pathway is activated as observed by an increase in their corresponding receptors and enzymes implicated. Furthermore, knockdown of the resolvins E and D receptors impairs the induction of senescence. This pathway is conserved not only during senescence but also in fibroblasts derived from aged human individuals, aged mice and during other inflammatory responses. A metabolomics analyses shows an increase in different precursors of resolvins in senescence. In accordance with prior data, we find that small extracellular vesicles (sEV) isolated from young human donors ameliorate inflammation and the biogenesis of resolvins both in different cell models and in aged mice. In summary, here we present data showing that the resolvins biogenesis pathway is induced in ageing and cellular senescence.

## INTRODUCTION

Ageing is a natural process by which a cellular, tissue and whole organism decline in functionality is observed. The process of ageing is mediated by different biological processes, one of which is the activation of cellular senescence and altered intercellular communication(Lopez-Otin et al., 2023). Senescence is a cellular phenotype characterized by a stable cell cycle arrest. More predominantly, senescent cells have a very active secretome termed the senescence-associated secretory phenotype (SASP) which mainly comprises cytokines, chemokines, metalloproteases and extracellular vesicles(Fafian-Labora and O’Loghlen, 2020; Mitsuhashi et al., 2013).

Resolvins are lipid mediators derived from the oxygenation of polyunsaturated fatty acids (PUFAs), such as the omega-3 fatty acid. They can be categorised into E-series or D-series resolvins. They are bioactive lipids which mediate signalling in an autocrine and paracrine manner through the binding to G protein-coupled receptors (GPCRs). They mediate a plethora of functions including inflammation, metabolism, cell signalling and senescence. Although other bioactive lipids have been implicated in senescence(Loo et al., 2017; Wiley et al., 2021), the role that resolvins play as part of the SASP is less characterized.

Here, we describe a role for a variety of resolvins (RvE1, RvD1 and RvD2) in regulating senescence both *in vitro* and in naturally aged mice. We have identified that the biosynthesis pathway for resolvins is activated upon the induction of senescence by expressing their respective binding receptors and the enzymes implicated in their biogenesis. This is consistent in senescence induced by various triggers, different cells models and in ageing (both in human and mouse). Importantly, genetic manipulation of their receptor prevents the induction of senescence. A metabolomics analyses further identifies different precursors of resolvins during senescence. In addition, treatment of aged mice or aged/senescent cells with small extracellular vesicles (sEV), which we previously identified to ameliorate senescence (Fafian-Labora et al., 2020), also prevent the biogenesis of resolvins. In summary, here show that the resolvins biogenesis pathway is induced during ageing taking advantage of human and mouse models and cellular senescence.

## RESULTS

### Resolvin biogenesis increases during cellular senescence

In order to identify a role for the resolvin biogenesis pathway during cellular senescence, we determined the mRNA expression levels of different resolvin receptors (**Figure 1A**). For this, we took advantage of human primary foreskin fibroblasts (HFFF2) treated with etoposide which we have shown to induce DNA damage-induced senescence (DDIS) (**Figure S1A)**(Borghesan et al., 2019; Fafian-Labora et al., 2020; Rapisarda et al., 2017). As shown in **Figure 1B**, the mRNA expression levels of the RvE1, RvD1 and RvD2 receptors (*ERV1/ChemR28*, *DRV1/GPR32* and *DRV2/GPR18* respectively), were upregulated during senescence. RvE1 has also been shown to bind the receptor BLT1/LTB4R1 and RvD1 the receptor ALX/FRP2. However, no detection or changes were observed for these receptors during DDIS (not shown). We further validated our findings in HFFF2 expressing the oncogene H-RAS^G12V^ (iRAS) in a tamoxifen (4OHT)- inducible vector which activates senescence and HFFF2 expressing a control vector (iC) (**Figure S1B**)(Borghesan et al., 2019; Fafian-Labora et al., 2020; Rapisarda et al., 2017). As the biogenesis of resolvins is induced upon the activation of specific enzymes, Cyclooxygenase (COX-2) and Lipoxygenase (5-LOX, 15-LOX), we next determined the expression levels of these enzymes during senescence (**Figure 1C**). In fact, we could observe an increase in the mRNA levels of *COX2*, *ALOX15* and *ALOX5AP* (ALOX5 activating protein) during DDIS (**Figure 1D**). Interestingly, we could also observe a positive correlation between the biomarkers of senescence, *CDKN2A* and *CDKN1A* and markers of resolvin biogenesis *ALOX15* and *ALOX5AP* (**Figure 1E**). In addition, the percentage of cells positive for 5-LOX by IF in human primary fibroblasts stably expressing H-Ras^G12V^ is also higher (**Figure 1F**). Thus, we show that there is an increase in the biogenesis of resolvins E1 and D1/2 during senescence.

**Figure 1.**
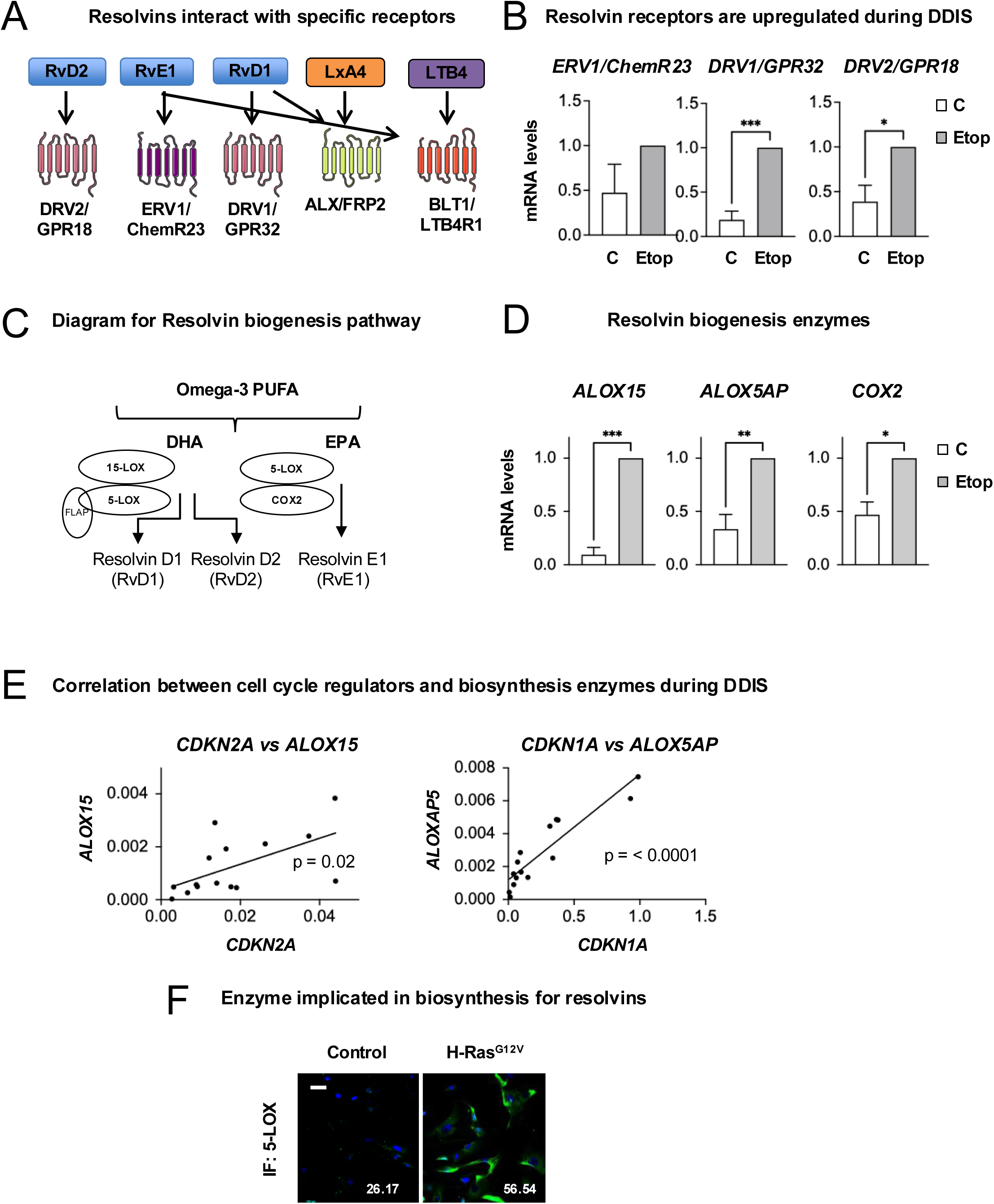
The resolvin biosynthesis pathway in activated during senescence. **(A)** Diagram showing which Resolvin (Rv) interacts with which receptor. RvE1 can bind either *ERV1/ChemR23* or *BLT1/LTB4R1* (Leukotriene B4 Receptor 1), while RvD1 can bind either *DRV1/GPR32* (G protein-coupled receptor 32) or *ALX/FRP2* (Formyl Peptide Receptor 2). RvD2 binds *DRV2/GPR18*. Other lipid mediators, LxA4 (LipoxinA4) and LTB4 (Leukotriene B4) also bind *ALX/FRP2* and *BLT1/LTB4R1* respectively. **(B)** Quantitative PCR analysis for HFFF2 cells undergoing DNA-damage induced senescence (DDIS) by treatment with Etoposide (Etop) 50μM for 7 days. The mRNA levels of different Resolvin receptors were determined: *ERV1*, *DRV2* and *DRV2*. **(C)** Diagram showing the biosynthesis pathway for resolvins series-D and -E from the omega-3 polyunsaturated fatty acids (PUFA): DHA (Docosahexaenoic Acid) and EPA (Eicosapentaenoic acid). The enzymes implicated are Lipoxygenases (5-LOX and 15-LOX), the 5-LOX activating protein (FLAP or ALOX5AP) and Cyclooxygenase 2 (COX2). **(D)** mRNA levels for enzymes implicated in the biogenesis of resolvins E and D in HFFF2 undergoing DDIS for 9 days. Graphs represent the mean ± SEM of 3 independent experiment. **(E)** Reanalysis of the data in (D) show a positive correlation between biomarkers of senescence (*CDKN1A*, *CDKN2A*) and resolvins biogenesis enzymes (*ALOX15*, *ALOX5AP*) in HFFF2 cells upon DDIS at different time points. Data represent the mean ± SEM of 3 independent experiments. **(F)** Representative image and quantification showing the expression levels of 5-LOX by IF in H-RAS^G12V^ cells. Data show the mean ± SEM of 3 independent experiment. See also **Figure S1**.

**Figure S1.**
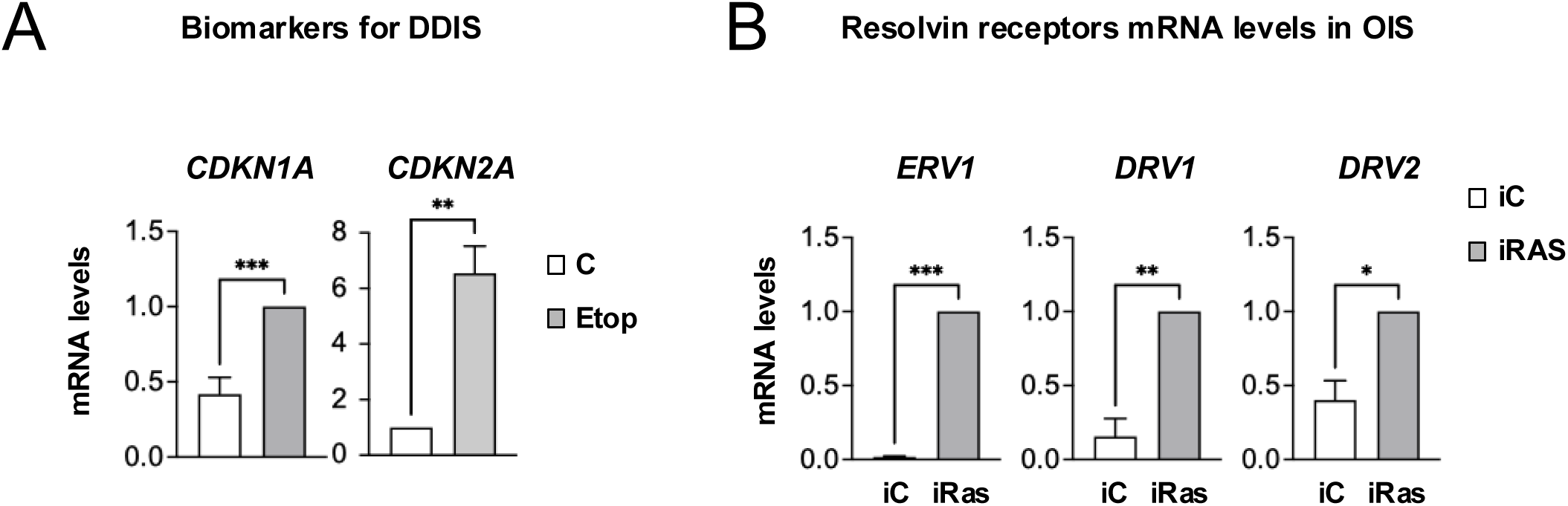
Resolvins are upregulated in cellular senescence in human primary fibroblasts cell cultures. Related to Figure 1. (**A**) mRNA levels quantification to confirm the induction of DDIS upon treatment with Etoposide (Etop) in HFFF2 fibroblasts. Data represent the mean ± SEM of 3-7 independent experiment. 1E (**B**) Quantitative PCR to evaluate the expression of the resolvin receptors during OIS (oncogene-induced senescence). Data represent the mean ± SEM of 3-5 independent experiment. Welch test analysis was performed. (A) is done by performing a reanalysis of the data from Figure 1E.

### Resolvin biogenesis is activated during ageing in mice and human primary fibroblasts

In order to validate our results during ageing, we took advantage of human primary fibroblasts isolated from healthy young (1-3 years old) and old (∼80 years old) donors (**Figure 2A**), which present several biomarkers of senescence (**Figure S2A**)(Fafian-Labora et al., 2020). In accordance with an activation of the resolvin biogenesis pathway during senescence, we could also confirm an increase in the mRNA expression levels of resolvins E and D receptors (**Figure 2B**) and their corresponding biogenesis enzymes (**Figure 2C**). Interestingly, in this model, we could determine an increase in the mRNA levels of *ALX* suggesting the RvD1 might also bind this receptor during ageing (**Figure S2B**), while no detection was observed for *BLT1* expression. To further validate our findings in ageing, we took young (∼2 months) and old (∼ 24 months) mice and quantified the expression levels for receptors and biogenesis enzymes in different tissues (**Figure 2D**). As expected, we could observe an increase in different biomarkers of senescence and ageing in the cortex such as *Cdkn2a* (**Figure S2C**) and *Il1a*, *Il1b* (**Figure S2D**). This increase is concomitant with an increase in *Erv1* and *Drv2* mRNA levels (*Drv1* does not exist in mice) (**Figure 2E**) and their respective biogenesis enzymes, *Alox5, Alox5ap* and *Cox2* (**Figure 2F**). We could also observe an increase in the expression levels of the biogenesis enzyme 5-LOX in the cortex of young and old mice by immunohistochemistry (IHC) (**Figure 2G**). Interestingly, *in vivo* we could detect an increase in the mRNA expression levels of *Blt1* in the cortex in aged mice with little changes observed in the striatum and hippocampus (**Figure S2E, F**). However, an increase in *Alx* mRNA expression levels in aged mice was observed in the cortex, hippocampus, striatum and spleen (**Figure S2G**). Altogether, our data show an increase in the genes necessary to induce an activation of the biogenesis of resolvins E1 and D1/2 in ageing in human and mouse samples.

**Figure 2.**
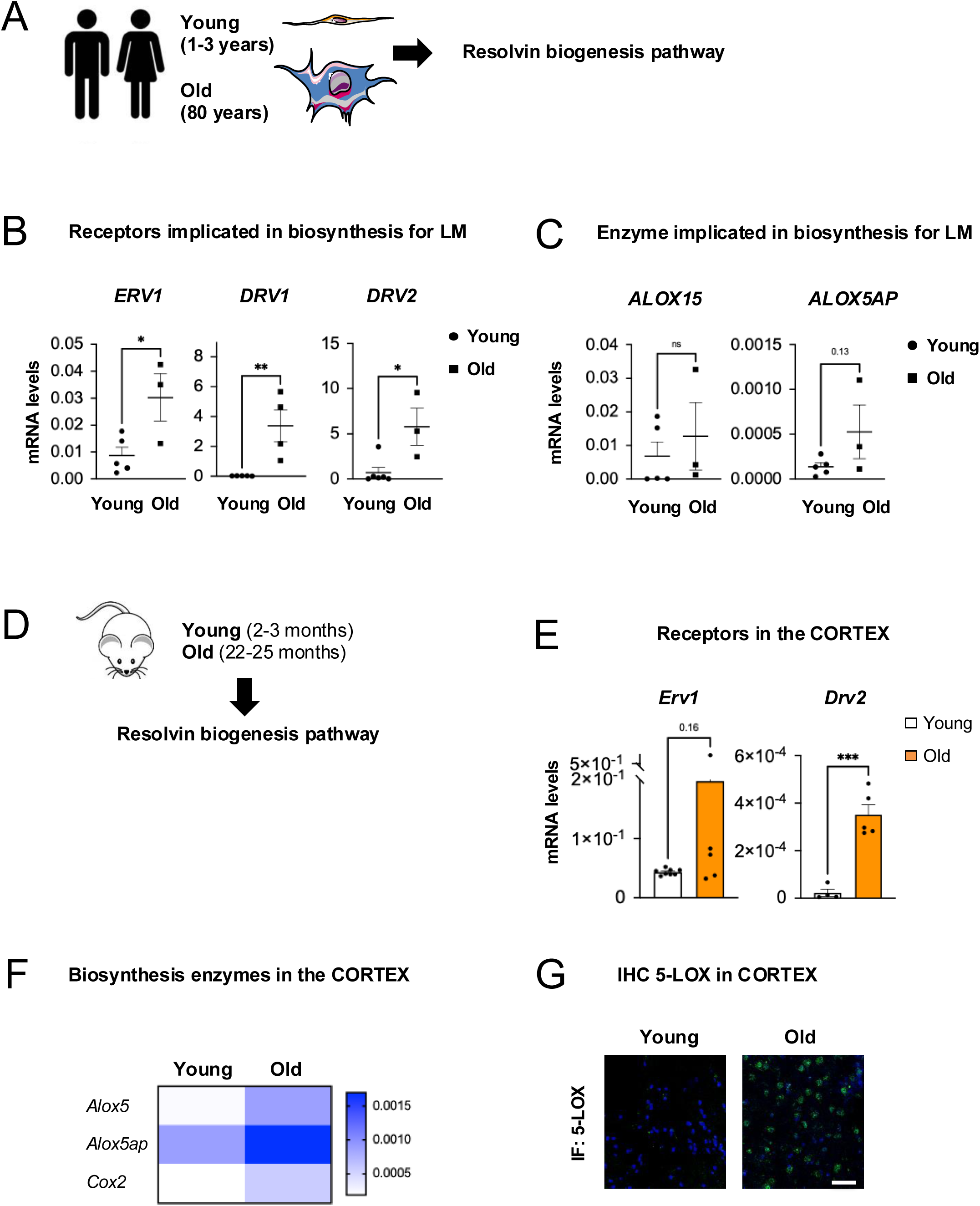
The resolvin biosynthesis pathway in activated during ageing in human and mouse. **(A)** Diagram of the experimental settings in human primary cell fibroblasts isolated from young (1-3 years) and old (approximately 80 years) donors. **(B)** *ERV1*, *DRV2* and *DRV2* mRNA levels were determined by qPCR in 5 young (GM05399, GM05565, GM02036, GM05758, GM00969) and 3-4 old human primary fibroblasts donors (AG06240, AG13222, AG16086, AG13152) cell cultures. **(C)** The mRNA expression levels for enzymes implicated in resolvin biogenesis was quantified. **(D)** Diagram of the experimental settings performed in young (2-3 months) and aged (22-25 months) mice. (**E, F**) Quantitative expression levels of the (**E**) resolvin receptors and (**F**) biogenesis enzymes in the cortex of young and old mice. (E, F) *Rps14* was used as housekeeping gene of reference. Graphs show the mean ± SEM of 4-8 young mice and 5 old mice. (**G**) Immunohistochemistry for the expression levels of 5-LOX in the cortex of young and old mice. Scale bar, 50μm. See also **Figure S2**.

**Figure S2.**
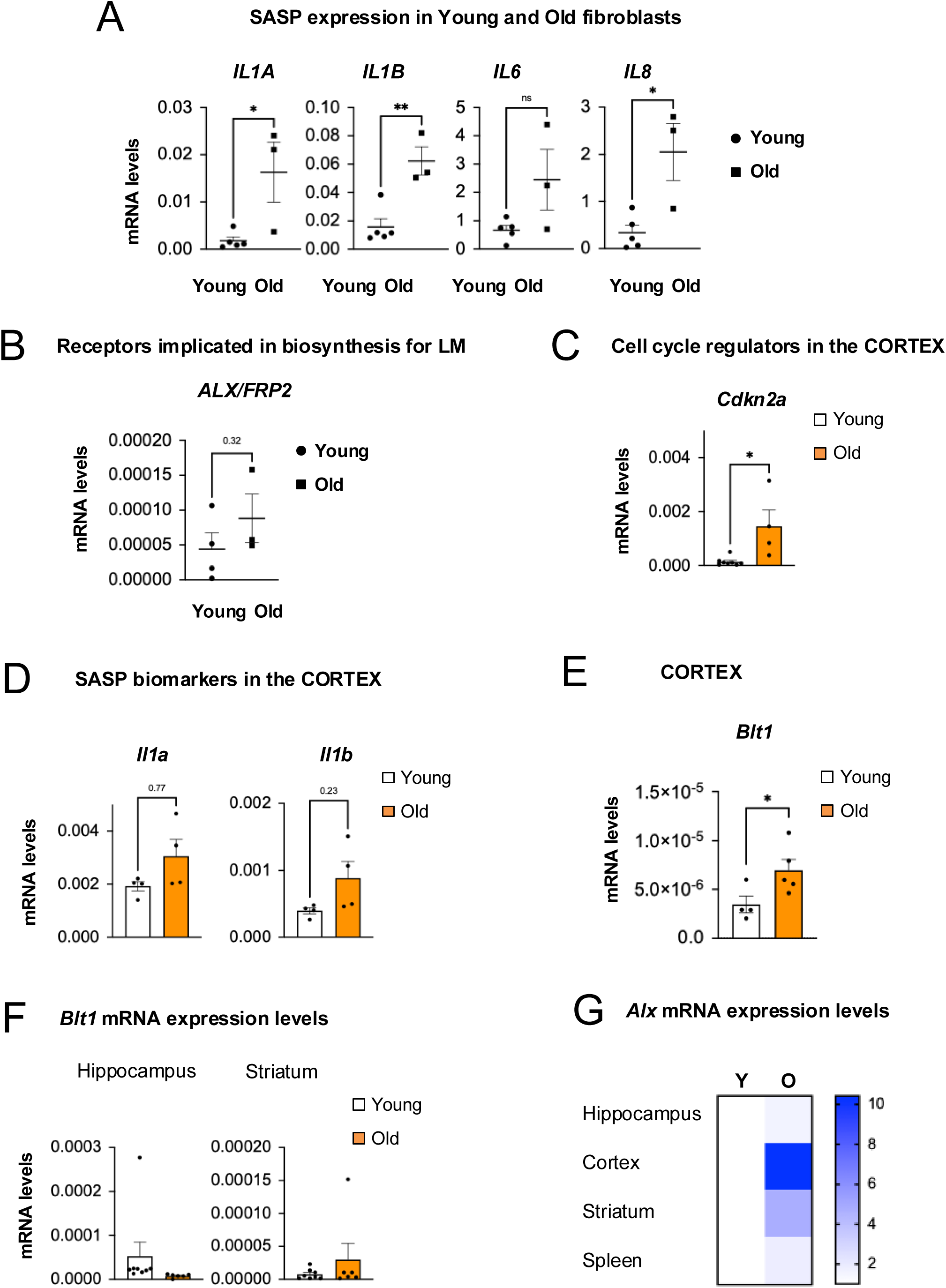
Resolvins are upregulated in ageing in human and mouse models. Related to Figure 2. (**A**) Human primary fibroblasts from old donors present several biomarkers of senescence(Fafian-Labora et al., 2020). (**B**) mRNA expression levels for receptors implicated in the biogenesis of resolvins in the same cell cultures as Figure 2. All data represent the mean ± SEM of 5 young and 3,4 old human primary fibroblasts. (**C, D**) qPCR data show an upregulation of (**C**) *Cdkn2a* and (**D**) SASP factors in the cortex of aged mice. Graphs show the mean ± SEM of 4-8 young and 4 old mice. (C, D, E) Data are represented using *Rps14* as housekeeing gene. (**E**, **F**) Expression levels for *Blt1* in the (**E**) cortex and (**F**) hippocampus and striatum in young and old mice. (**G**) qPCR analyses for *Alx* expression in young and old mice in different tissues. Graphs show the mean ± SEM of 4 young mice and 4 old mice.

### Resolvin biogenesis is also increased in brain microglia

As we observe an increase in the resolvin biogenesis pathway during senescence and ageing in the mouse brain (**Figure 2**), we next decided to determine whether this pathway is activated in microglia isolated from mice (**Figure 3A**). To induce senescence and try to mimic a chronic inflammatory state, we treated isolated brain microglia with 5 μg/ml of Lipopolysaccharide (LPS) for 48h, washed the cells and collected after 5 days for analysis (**Figure 3B**). In fact, after LPS treatment we could observe an increase in different biomarkers for senescence (**Figure 3C)**, an increase in the release of Il1b measured by ELISA (**Figure 3D**) and significant changes in morphology (**Figure 3E**). Concomitant with an increase in different biomarkers of senescence, we could also observe an increase in the expression levels of the receptor *Erv1* (**Figure 3F**), further supporting the hypothesis that the resolvin pathway is activated during senescence in different models.

**Figure 3.**
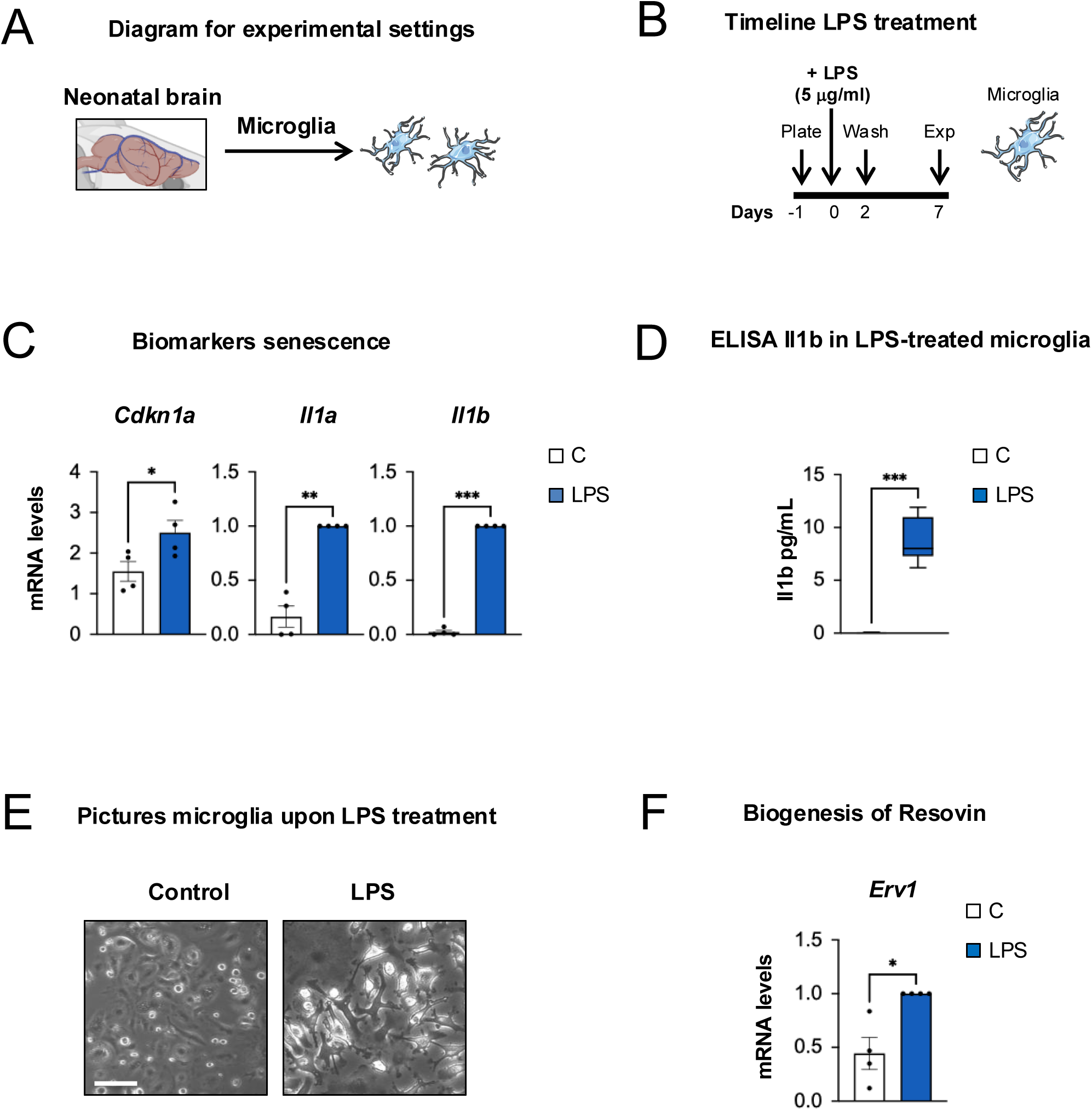
Resolvin biogenesis correlates with senescence in LPS-treated brain microglia. **(A)** Microglia from neonatal mice (P1-P5) were isolated and **(B)** treated with LPS (5 μg/ml) for 2 days following a 5 day rest to mimic chronic inflammation. **(C)** qPCR data show upregulation of different senescence biomarkers upon LPS treatment. Data represent the mean ± SEM of 4 independent experiments. **(D)** ELISA for Il1β upon LPS treatment in brain microglia show an increase in the levels of Il1β in the conditioned media. (**E**) Representative pictures showing the morphological changes of brain microglia upon LPS treatment. Scale bar, 50μm. (**F**) Concomitant with an induction in different biomarkers of senescence we observe an induction in the resolvin pathway shown by measuring the mRNA levels of *Erv1* upon LPS treatment.

### Functional implication of the resolvin biogenesis pathway in the activation of cellular senescence

To confirm a functional role for the resolvin biogenesis pathway during senescence, we next generated shRNA targeting *DRV1* and *ERV1* in IMR90 fibroblasts and analysed the expression of different biomarkers of senescence. We can actually see that there is a partial prevention in the upregulation of biomarkers of senescence such as *CDKN2A* (**Figure 4A**) and components of the SASP (**Figure 4B**) upon shDRV1 and shERV1. As per our preliminary data, we also see a partial prevention of enzymes implicated in the biogenesis of resolvin upon *DRV1* and *ERV1* knockdown (**Figure 4C, D**). These data are further validated at the level of SASP released to the conditioned media by taking advantage of a Human Cytokine Array (**Figure 4E, F**). Altogether, our data show that the knockdown of the resolvin receptors *ERV1* and *DRV1* partially prevents the induction of senescence and the increase in the activity of the resolvin biogenesis pathway during senescence.

**Figure 4.**
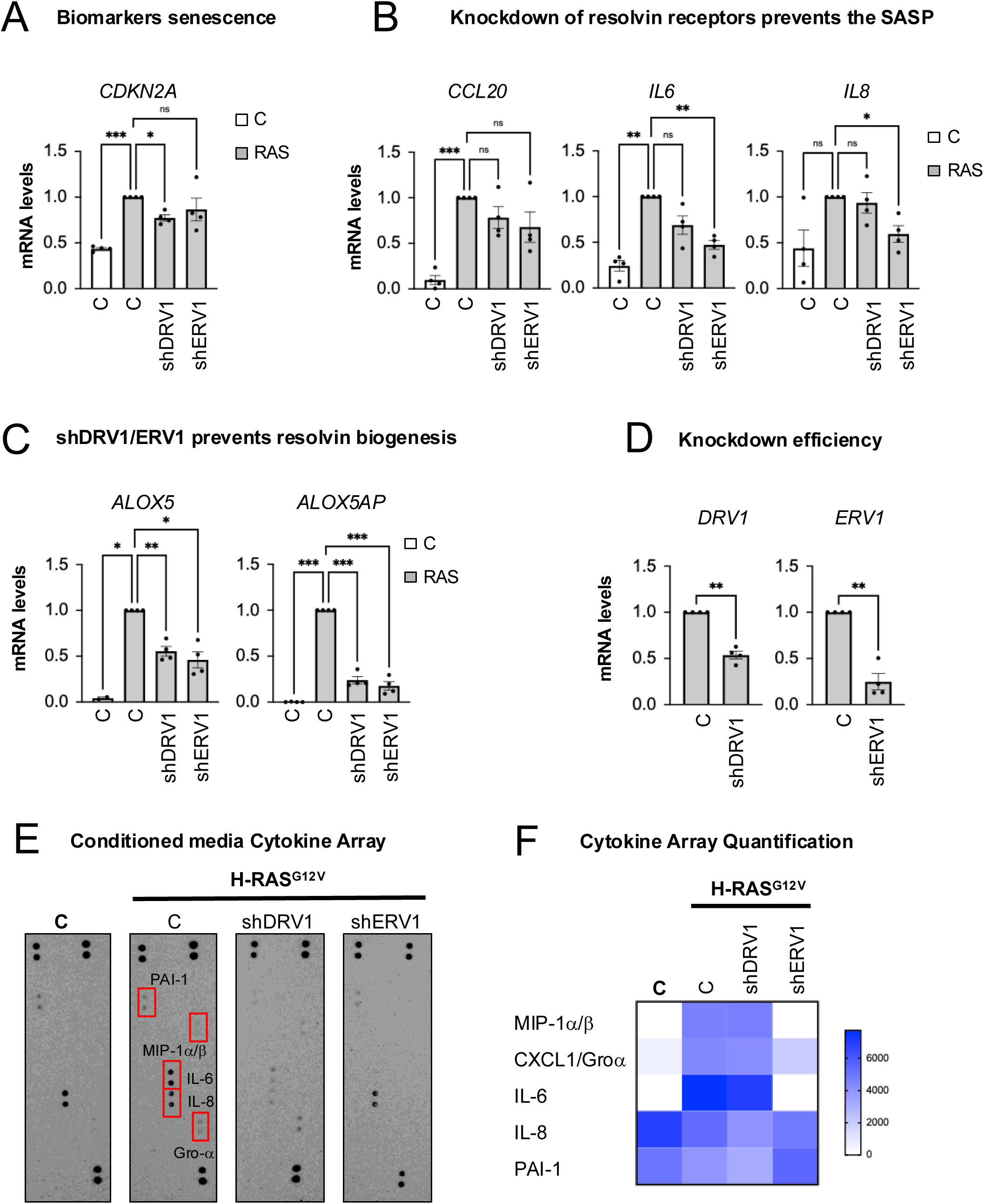
Knockdown of the resolvins receptors partially prevent the induction of senescence. (**A, B**) Knockdown of *DRV1* (shDRV1) and *ERV1* (shERV1) partially prevents the upregulation of (**A**) *CDKN2A* and (**B**) several SASP factors, concomitant with a reduction in the expression levels of different (**C**, **D**) resolvin biogenesis enzymes, *ALOX5* and *ALOX5AP*. (**E, F**) Human cytokine array and quantification shows a decrease in several SASP factors released to the conditioned media upon shERV1 and shDRV1.

### Lipid mediator targeted metabolomics analysis in senescent cells

The biogenesis of resolvins can derive from an increase in the abundance in polyunsaturated fatty acids (PUFA). Thus, to investigate whether PUFAs are responsible for the increase in the biogenesis of RvE1 and RvD1/2 during senescence, we performed a targeted lipid mediator metabolomics analyses taking advantage of LC-MS/MS in Control and H-Ras^G12V^ fibroblasts (**Figure 5A, Figure S3A**). This approach identified 295 peaks which were detected and annotated according to the Human Metabolome Technology (HMT) standard library. To identify differences between groups an OPLS-DA plot was performed. In fact, **Figure 5B** shows that the lipid mediator composition from control and H-Ras^G12V^ fibroblasts are significant drivers for group differences. In addition, a correlation table (**Figure 5C**) shows a linear correlation between control and senescent samples, while hierarchical clustering of the top 50 mediators identified show differential clustering between samples (**Figure 5D**). Furthermore, we observe an increase in the mean of the resolvins E and D precursors, DPA (Docosapentaenoic acid), DHA (docosahexaenoic acid), and EPA (eicosapentaenoic acid) in the H-Ras^G12V^ sample (**Figure 5E**). Altogether, our data show that there is an increase in the fatty acid lipid mediator precursors of resolvins D and E in senescence.

**Figure 5.**
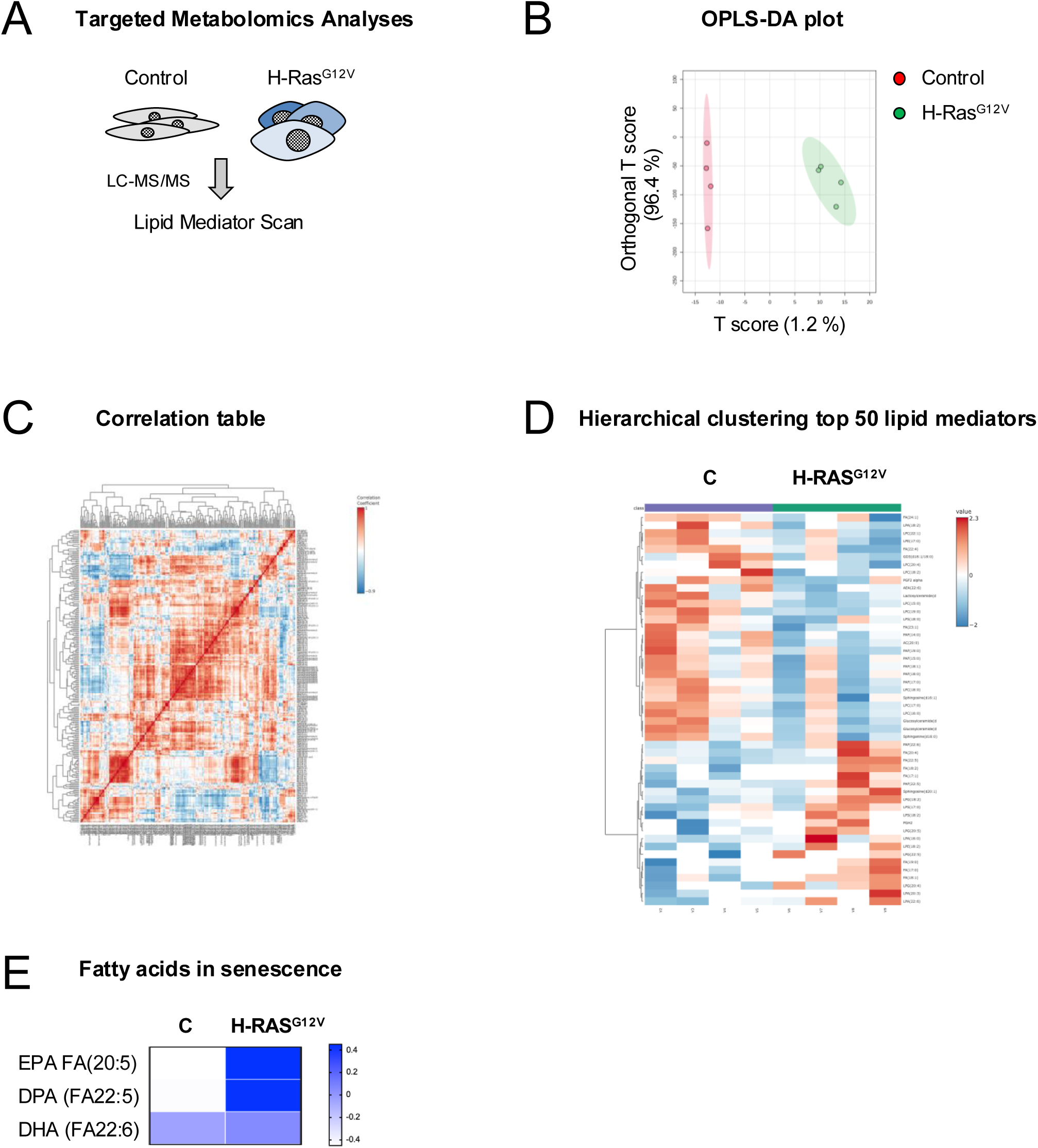
Targeted lipid mediator analyses in H-Ras^G12V^ expressing IMR90 fibroblasts. (**A**) Schematic representation of the lipid mediator metabolomics analysis. (**B**) OPLS-DA plot between control and H-Ras^G12V^ cells show significant differences between both groups. (**C**) Linear correlation between control and H-Ras^G12V^ samples. (**D**) Hierarchical clustering of the top 50 mediators identified by t-test analyses. (**E**) Heatmap of the mean of the precursors for resolvins series -D and -E in control and H-Ras^G12V^ samples. DPA (Docosapentaenoic acid), DHA (docosahexaenoic acid), and EPA (eicosapentaenoic acid). All samples are the representation of 4 biological replicates per group. Related to **Figure S3**.

**Figure S3.**
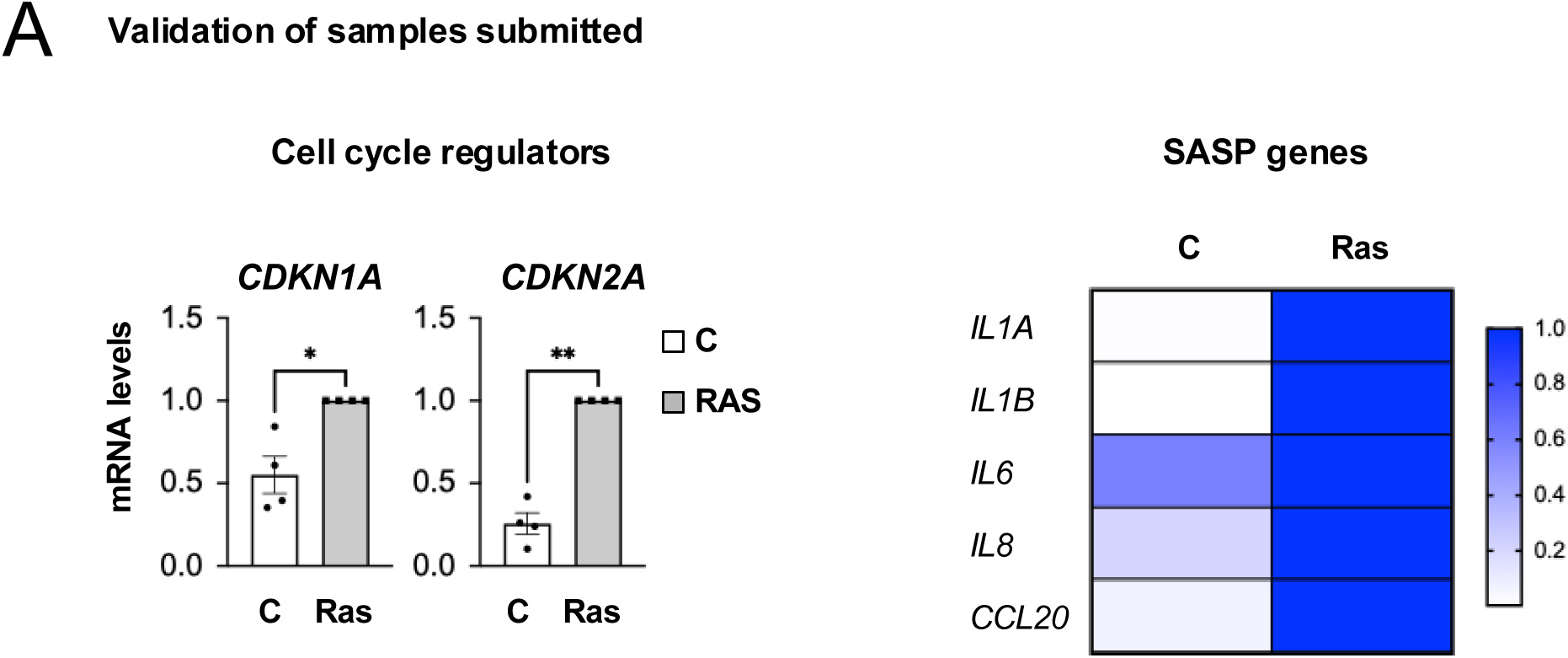
Validation of senescence in the samples submitted for Lipid Mediator metabolomics profiling. (**A**) qPCR analysis for different biomarkers of senescence of the samples submitted for metabolomics. Data show the mean of 4 biological replicates. **Related to** Figure 5.

### Small Extracellular Vesicles from young individuals reduce the biogenesis of resolvins during senescence and ageing

Previous data from our lab show that small extracellular vesicles (sEV) from young individuals reduce the burden of senescence(Fafian-Labora et al., 2020) (**Figure S4A**). To determine whether sEV isolated form human primary fibroblasts from young individuals also ameliorate the increase observed in the biogenesis of resolvins during ageing, we isolated sEV by serial ultracentrifugation as previously(Borghesan et al., 2019; Fafian-Labora and O’Loghlen, 2021; Fafian-Labora et al., 2020; Menendez-Garcia M, 2025 ). For this, we took advantage of young and aged mice and treated them via intraperitoneal injection (IP) with sEV for 3 weeks as before (**Figure 6A**)(Fafian-Labora et al., 2020). Interestingly, concomitant with an amelioration in different senescence biomarkers as observed before (**Figure S4A**)(Fafian-Labora et al., 2020), we could observe a reduction in the resolvin biogenesis pathway in different tissues such as the spleen and different regions of the brain (**Figure 6B-D**). Altogether, these data show that the resolvin biogenesis pathway is ameliorated concomitantly with senescence during ageing in human and mouse models.

**Figure 6.**
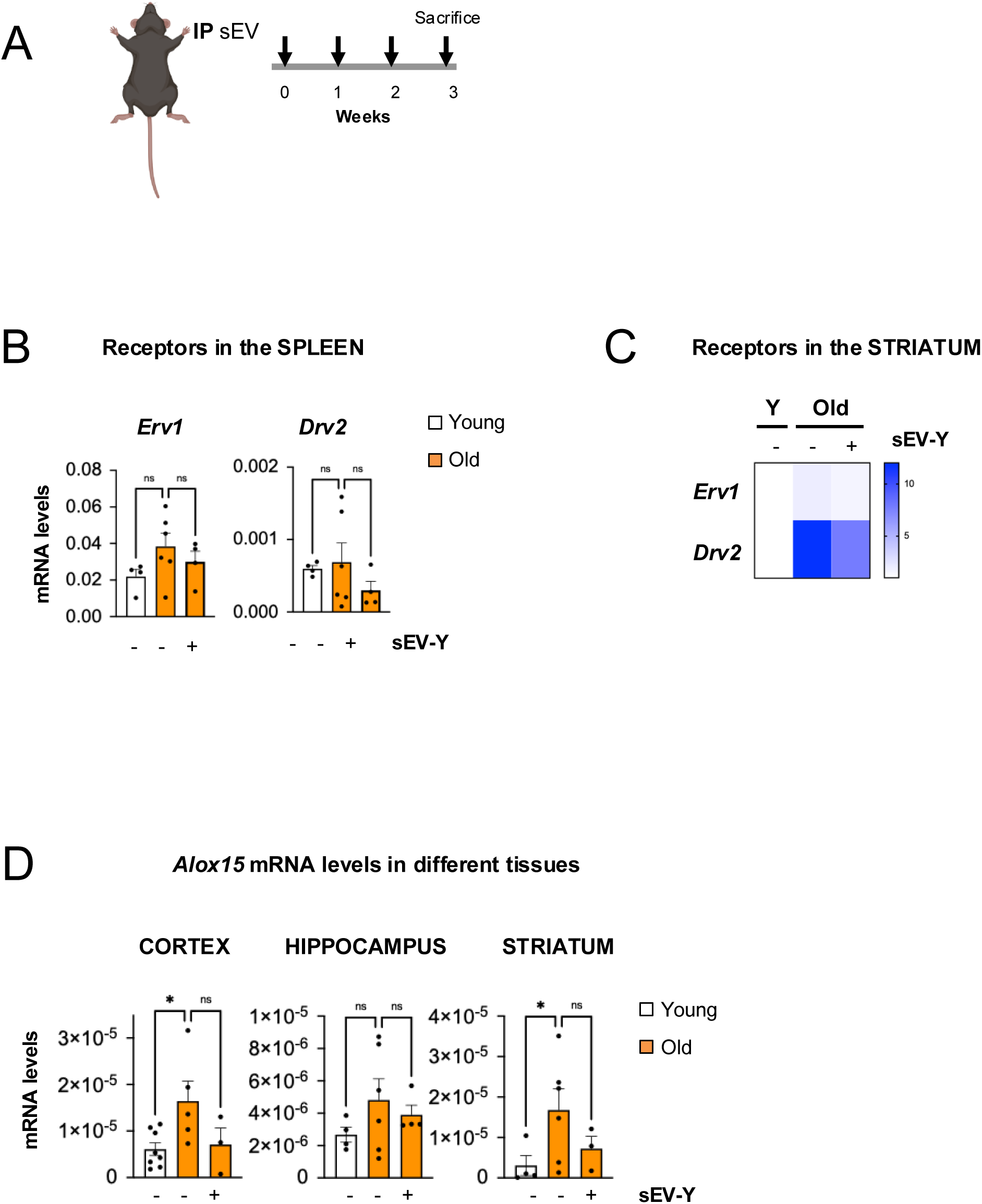
The resolvin biogenesis pathway is downregulated in aged mice by sEV treatment. (**A**) Schematic representation of the repeated treatments of sEV in aged mice. (**B-D**) qPCR analyses show upregulation of different (**B**, **C**) resolvin receptors and **(D)** their biogenesis enzymes in different tissues of aged mice. Data show the mean ± SEM of 3-8 mice per condition. Data normalised to *Rps14*. **Related to Figure S4**.

**Figure S4.**
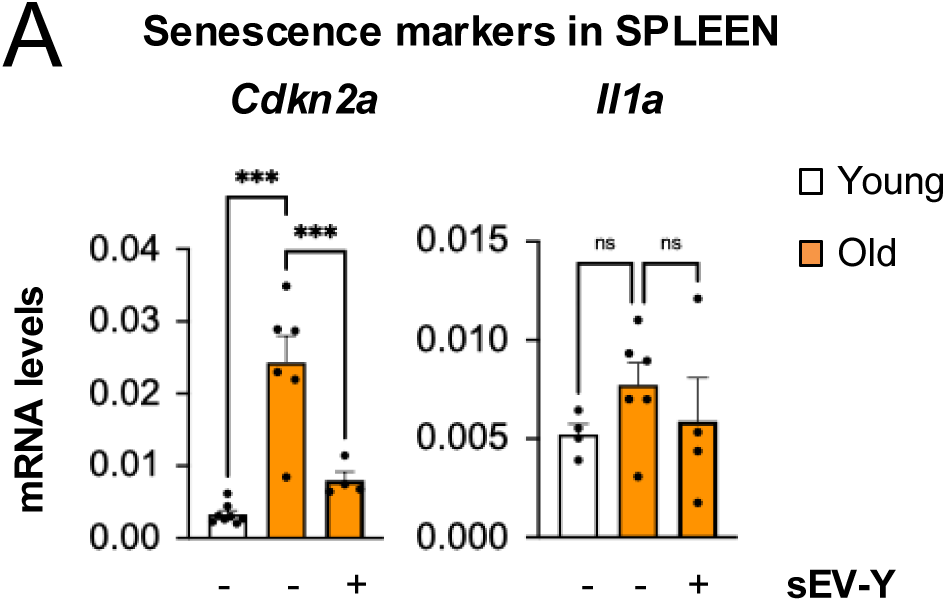
Validation of diverse biomarkers of senescence upon sEV treatment in aged mice. (**A**) qPCR analysis of *Cdkn2a* and *Il1a* in the spleen of young and aged mice treated with sEV. **Related to** Figure 6.

Next, wanted to determine whether this was also conserved in human primary cell cultures. Thus, we treated iRas cells with sEV and determined the mRNA expression levels of *ERV1*, *DRV1/2*. As show in **Figure 7A-B**, treatment of iRAS with sEV reduced the increased expression of *ERV1* and *DRV1* receptors (**Figure 7B**). A similar trend was observed when treating different old donors with sEV (**Figure 7C-D**). Thus, sEV have the ability to reduce the biogenesis of resolvins induced during ageing and senescence.

**Figure 7.**
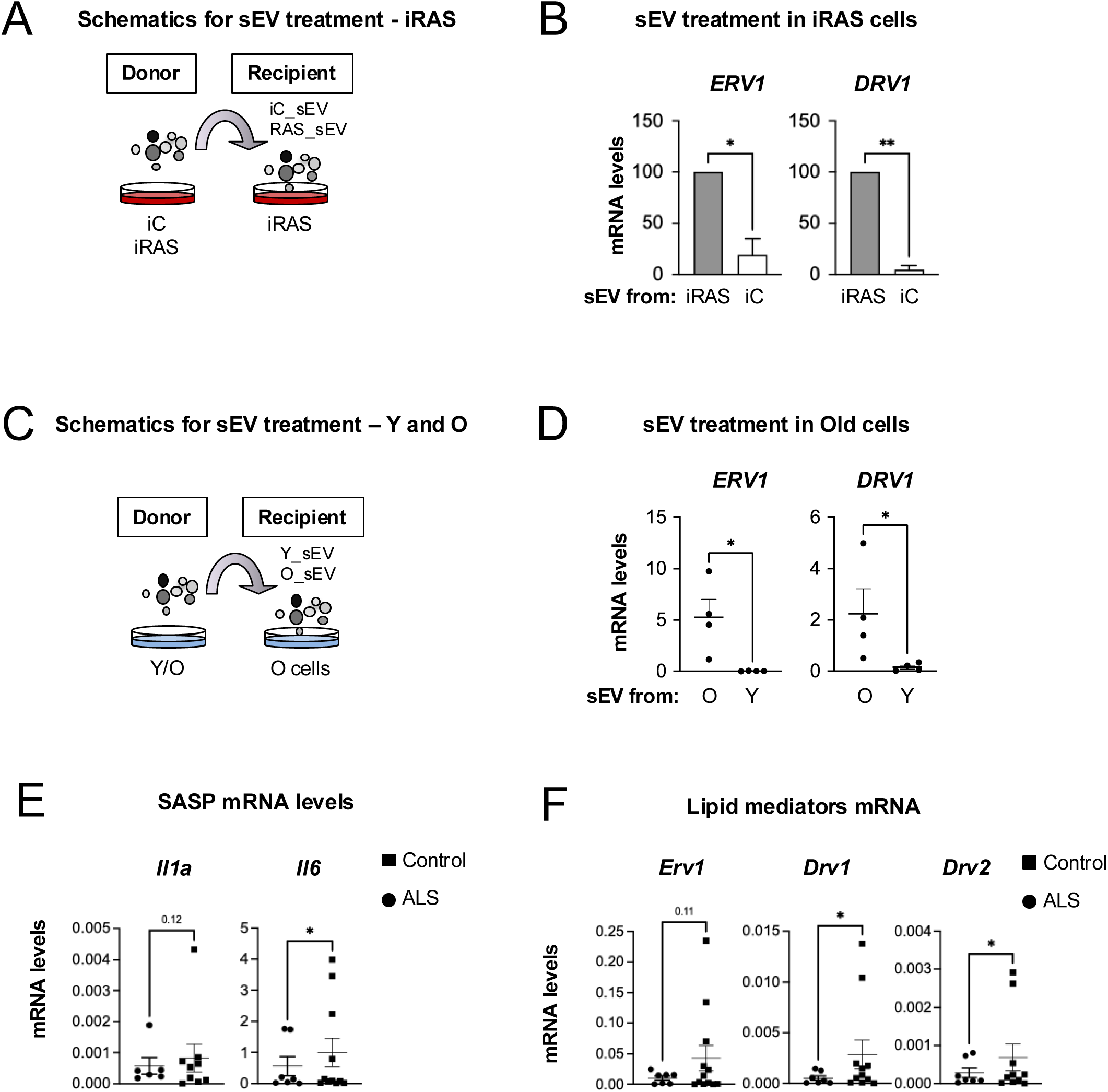
Small Extracellular Vesicles prevents the increase in the resolvin biogenesis pathway in ageing. **(A)** Diagram showing the experimental settings in which iRAS cells are treated with sEV from either iC or iRAS cells. **(B)** qPCR analysis for the receptors *ERV1*, *DRV1* mRNA was performed. **(C)** Schematic representation of the experimental settings in which Old human primary fibroblasts are treated with sEV isolated from either Young or Old cells. **(D)** qPCR analysis for the resolvin receptors was performed. Data represent the mean ± SEM of 4 young (GM05399, GM00969, GM05565, GM05758) and 4 old (AG16086, AG06240, AG13152, AG13222) donors. **(E)** mRNA levels for different SASP factors in ALS-derived fibroblasts correlates with an (**F**) increase in the expression levels of the resolvin receptors *ERV1*, *DRV1*, *DRV2*. Graphs show the data for the mean ± SEM of 6 healthy controls and 9-12 ALS patient-derived fibroblasts.

### The resolvin biogenesis pathway is enriched in ALS-derived fibroblasts

Ageing and age-related diseases present a highly prominent inflammatory profile. Amyotrophic Lateral Sclerosis (ALS) is a motoneuron disease with a prominent and systemic inflammatory response. As previous data have shown an increase in *DRV1* and *DRV2* levels in blood associated with increase ALS survival (Yildiz et al., 2025), we set to evaluate the activation of the resolvin biogenesis pathway in fibroblasts isolated from ALS patients in comparison with healthy controls. As expected, ALS-derived fibroblasts presented high levels of inflammatory cytokines (**Figure 7E**), in accordance with an induction of cellular senescence and ALS disease. Furthermore, this correlated with an increase in the mRNA expression levels of *ERV1*, *DRV1* and *DRV2* (**Figure 7F**), suggesting a correlation between the induction of senescence and the increase in the biogenesis of resolvins. Thus, the resolvin pathway is also activated in age-related inflammatory diseases such as ALS.

## DISCUSSION

The importance of the presence of senescent cells during ageing and in age-related diseases is becoming more and more prominent. Thus, understanding the mechanisms by which these cells contribute to the tissue microenvironment is crucial. The intercellular communication of senescent cells is extremely varied, from cell-to-cell contact, fusion, cytoplasmic bridges, ECM interactions and others(Fafian-Labora and O’Loghlen, 2020). However, the better characterised means of communication is through the SASP. In spite of this, most studies focus exclusively in cytokines and chemokines as part of the SASP, while the contribution of EV and metabolites remains less characterised(Fafian-Labora and O’Loghlen, 2020).

The contribution of senescent cells to the microenvironment is paradoxical and confusing. On the one hand, they can be proinflammatory, but on the other hand they have been described to be anti-inflammatory. As resolvins play a key role in tissue resolution as anti-inflammatory signalling molecules, here we set to determine whether the resolvin biogenesis pathway was implicated in senescence. Our results show that not only the resolvin biogenesis pathway is activated during senescence in different models but also in human and mouse ageing and age-related diseases. These findings are interesting, as senescent cells are known to recruit immune cells in different scenarios *in vivo* highlighting the potential of senescence in tissue resolution(Wang et al., 2024). Our results are consistent with previous findings, where changes in the lipid composition of senescent cells was described(Quijano et al., 2012) and an increase in the abundance of EPA and DHA has been shown in different models of senescence(Wiley et al., 2021). Furthermore, changes in the release of alternative lipid mediators such as prostaglandins(Loo et al., 2017) or in the expression levels of the biogenesis enzymes(Catalano et al., 2005; Goncalves et al., 2021) has also been shown in senescence independently, highlighting the complexity of senescence in its pro- and anti-inflammatory contribution (Loo et al., 2017; Wiley et al., 2021).

In contrast with OIS and DDIS, during human and mouse ageing we do see an increase in the expression levels of the receptor *ALX*. ALX is bound not only by RvD1 but also by lipoxin A4 (LxA4) suggesting that the regulation of LxA4 could be something specific to ageing. However, the fact the mice lack DRV1 could also indicate that Alx is compensating for the lack of DRV1 during ageing in mice. Interesting, we found an increase in the expression of *Blt1* only in the cortex of aged mice. As it is primarily expressed in granulocytes, monocytes and lymphocytes, this could indicate that there is an increase in the infiltration of immune cells in the cortex of aged mice during ageing.

In summary, here we identify that resolvin biogenesis is increased in senescence and ageing in different cellular models and naturally aged mice and open the window to explore senescence as a pro-resolving cellular state.

## ACKNOWLEDGEMENTS

We thank the Coriell Institute for providing human primary fibroblasts from young and old donors. MMG received a PhD fellowship from Comunidad de Madrid (PIPF-2023/SAL-GL-29915). AO’s lab is supported by projects CNS2022-135134 financed by MICIU/AEI/10.13039/501100011033 and by the EU NextGenerationEU/PRTR, project PID2021-125656OB-I00 financed by MICIU/AEI/10.13039/501100011033/10.13039/501100011033 and by FEDER, UE and project SenesceX S2022/BMD-7393 financed by the Comunidad de Madrid.

## AUTHOR CONTRIBUTIONS

AO conceived and designed the study. MMC, MMG, AMN, CPL performed most of the experiments. CAB and JM performed the microglia experiments. IP and AM provided advise on the senescence and ALS experiments respectively. AO wrote the manuscript, with input from all the authors.

## STAR METHODS

### Mice treatments

Male C57BL/6 mice aged 3 months for young and 22-25 month-old mice were used for this study. Intraperitoneal injections (IP) were performed once a week to inject either 10 µg of isolated sEV or an equivalent volume of PBS for 3 weeks. Mice were sacrificed 24 h after the last injection. Tissues were divided in sections and stored adequately according to the analysis to be performed. All procedures used in animal experiments followed Spanish (RD 53/2013) and European legislation (2010/63/EU) and were previously approved by the CSIC and Madrid Region Committee (Spain).

### Cell culture, retroviral and lentiviral infections

IMR90 and HFFF2 fibroblasts were obtained by the Culture Collections (Public Health England, UK). Young and old primary human fibroblasts were obtained from the Coriell Cell Repository with the following codes: Young fibroblasts (GM05399, GM05565, GM02036, GM05758, GM00969) and Old fibroblasts (AG06240, AG13222, AG16086, AG13152). Cells were maintained in high-glucose, pyruvate, Dulbecco’s modified Eagle’s medium with 10% fetal bovine serum and 1% antibiotic-antimycotic solution. All cells were washed twice with PBS to remove excess FBS media and maintained in sEV-depleted FBS media for the duration of the experiment. FBS was depleted of sEV by overnight (ON) ultracentrifugation at 100,000 g at 4C (Optima L-100 XP Ultracentrifuge). The supernatant was removed and stored in 50mL falcons at -20C until required. CM was collected after 72h incubation with cells in 0,5% EV depleted FBS, unless specified otherwise. Brain microglia were cultured in DMEM-F12 supplemented with 10% FBS and 5 ng/mL GM-CSF. Methods used for retrovirus and lentivirus production and infection have been previously described (Acosta et al., 2008; Rapisarda et al., 2017).

### Microglia isolation and treatments

Mixed glial cultures were prepared from postnatal day (P)0–P5 mouse brains (C57BL6J). Pups were sacrificed by rapid decapitation, the brains were removed, and meninges were carefully peeled off. Cortices were mechanically dissociated by gentle trituration and the resulting suspension was filtered through a pre-wetted 70 µm cell strainer. Cells were centrifuged at 1200 rpm for 5min and cultured in DMEM-F12 (11320033, Thermofisher) supplemented with 10% FBS and 5 ng/mL GM-CSF (130-095-746, Milteny). Cultures were maintained at 37°C in 5% CO₂, with weekly medium changes. After 14 days *in vitro*, microglia were purified by orbital shaking of confluent mixed glial cultures at 150 rpm for 3–4 h, followed by collection of the supernatant and centrifugation. The microglial pellet was resuspended, counted, and plated onto poly-L-lysine–coated cultureware at 30,000–40,000 cells/cm² in complete medium containing GM-CSF, and allowed to recover for at least 48h prior to stimulation. Microglia purity (>99%) was tested by Iba-1 immunostaining. For stimulation, 5 ug/mL of lipopolysaccharide (E. Coli O26:B6 serotype, L8274, Sigma) was added to the media for 48h. Then, media were removed, cells were washed, and fresh medium was replenished. After 5 days, cells and conditioned media were harvested for downstream applications.

### Human Metabolon Metabolomic Lipid Mediator Analyses

Samples were submitted and processed by Human Metabolome Technologies (HMT), America, as follows:

#### Sample Preparation

The sample was mixed with 800 µL of 0.1% (v/v) formic acid in methanol containing internal standards by ultrasonication on ice. The sample suspension was then mixed with 550 µL of 0.1% formic acid in water. After the mixture solution was centrifuged (8,000 rpm, 4 C, 5 min), 800 µL of supernatant was mixed with 2,400 µL of 0.1% formic acid in water in a new 15 mL tube, and then purified by centrifugation (2,000 × g, 4C, 5 min) using SPE column (MonoSpinC18, 5010-2170, GL Sciences Inc., Tokyo, Japan). After the column was washed (centrifuged at 2,000 × g, 4C, 5 min) with 300 µL of 0.1% formic acid in water and 200 µL of 0.1% formic acid-25% methanol solution, lipid metabolites were eluted (centrifuged at 2,000 × g, 4□C, 5 min) with 200 µL of 0.1% formic acid in methanol.

#### Measurement

The compounds were measured in the Positive and Negative modes of LC-MS/MS based metabolome analysis in the following device: LC system: Agilent 1260 Infinity II + Agilent 1290 Infinity II High Speed Pump (Agilent Technologies Inc.). The column used was: Acquity UPLC HSS T3, 2.1 × 100 mm, 1.8 μm (Waters) and the MS system: AB Sciex QTRAP 5500 (AB Sciex).

#### Putative metabolites

From the LC-MS/MS measurement, 295 peaks were detected and annotated according to HMT’s standard library. Peak ID consisted of analysis mode and number where the measurement modes are as following: Cation (C), Anion (A), Positive (P) and Negative (N) mode. All putative metabolites were assigned based on m/z and MT or RT. Those listed in “PubChem ID / HMDB ID / peptide” were assigned on the basis of m/z only.

#### Data analyses

Data analysis was performed were performed using a statistical analysis software developed at HMT or the freely available software MetaboAnalyst.

### Human Cytokine Array and quantification

To measure changes in cytokine release upon ERV1 and DRV1 knock-down in senescent cells, the Human Cytokine Array from R&D Systems was used (#ARY005B). The protocol was carried out according to manufacturer’s instructions, using 0,7 ml of conditioned media for each experimental condition. Membranes were imaged using Chemidoc. Pixel density was measured using ImageJ (FIJI) software for each pair of duplicate spots representing each cytokine. The averaged background signal from each spot was subtracted and final signal results were compared between samples to determine the relative change for each cytokine.

### qPCR analysis

RNA was isolated using TRIzol Reagent or the RNeasy Mini Kit (QIAGEN, 74104) from the cells previously washed with PBS. Tissues were stored in TRIzol and frozen until further use. cDNA was generated using the High-Capacity cDNA Reverse Transcription Kit according the manufactureŕs instructions. qPCR was performed using SYBR Green PCR Master Mix on a 7500 Fast System RealTime PCR cycler. Primer sequences are listed in **Table S2**. The relative expression was calculated using the ΔΔCt methods using the Ct values generated using the 7500 software version 2.0.6. The data were normalized to a housekeeping gene, RPS14.

### Differential ultracentrifugation of several fractions from conditioned medium and sEV isolation

To isolate the EV fractions, the protocol of differential ultracentrifugation (Théry et al., 2006, 2018) was modified and adapted (Borghesan et al., 2019). All cells were plated for each individual experiment. Briefly, 1x10^6^ early passage donor cells were plated in a 10cm dish (10ml media) for each individual experiment. After 72h we isolated sEV. Briefly, the CM was collected, centrifuged at low speed (2,000 g for 20min; k-factor of the rotor is 41056) to eliminate dead cells and cellular debris prior to use. Large EV (lEV) were discarded after the 10,000g centrifugation step for 1h, washed in 15 ml PBS and spun down again at 10,000 g (k-factor of the rotor is 669) for 1h. The media was then filtered through a 0.22µm filter prior to the 100,000g centrifugation step. The sEV fraction was collected after a 1h and 20min 100,000 g centrifugation step and concentrated using a 10K column (Amicon Ultra-0.5 Filter) at 14,000 g for 10min obtaining a concentration factor 10X. The final 100,000 g pellet (sEV) was collected after a 1h and 20min and was washed once in 15ml of PBS and resuspended in 50 μl of PBS. An Optima L-100 XP Ultracentrifuge, with a Beckmann Fixed Angle Type 45 Ti rotor was used for sEV isolations. The k-factor of the rotor is 133.

### Immunofluorescence staining in cells

The cells were washed with PBS and fixed in 4 % (v/v) paraformaldehyde for 10-15 min at room temperature (RT). Following fixation, cells were washed in PBS and permeabilized with 0.4 % (v/v) Triton X-100 in PBS for 10 min at RT. Non-specific binding sites were blocked by incubating the preparations with 1x Blocking Solution (0.5% (m/v) BSA and 0.2 % (m/v) gelatin fish) for 1h at RT. Cells were incubated with the suitable concentration of primary antibody overnight at 4C. The next day, the preparations were washed three times with the Blocking Solution and incubated 1 hour with the appropriate secondary antibody. Subsequently, nuclei were counterstained with DAPI (0.3mg/ml) for 3 min. Finally, the cells were washed twice with 1x Blocking Solution and twice with PBS. The IF preparations were maintained with PBS and stored at 4C until the acquisition of pictures using a Confocal Laser Scanning Microscope (CLSM) LEICA TCS SP8 STED 3X.

### Immunofluorescence staining in tissues

Tissue sections on slides were fixed in paraformaldehyde at 4% for 30 minutes at room temperature. Then, they were washed 3 times with PBS and, afterwards, we performed heat-induced epitope retrieval in order to restore their antigenicity after fixation: Slides were submerged in 500mL of Tris-EDTA buffer solution (10mM Tris base, 1mM EDTA solution, 0.05% Tween 20, pH 9.0) and microwaved for 5 minutes twice, adding distilled water in between to replace the amount of evaporated buffer solution. Once the bath cooled down at room temperature, slides were washed twice with PBS and sections were blocked and permeabilised for 1 hour at room temperature with blocking buffer solution (5% bovine serum albumin, 5% normal goat serum, 0.5% Triton X-100, in PBS) in a humidity chamber (used for the remaining incubation steps in this protocol). Next, sections were incubated overnight with primary antibody at optimized concentration in blocking buffer solution at 4°C, washed 4 times with permeabilization buffer solution (0.1% Triton X-100 in PBS) and incubated for 1 hour with secondary antibodies diluted 1:500 in blocking buffer solution - from this step onwards, slides were kept in darkness. Then, sections were permeabilised 4 times for 5 minutes with permeabilization buffer solution and washed with PBS for 5 minutes. Finally, sections were incubated for 10 minutes with DAPI (0.3 µg/mL in PBS), washed for 5 minutes with PBS and coverslips were mounted on slides mounting solution and sealed with nail varnish. The acquisition of pictures was performed using a LEICA TCS SP8 STED 3X confocal microscope.

### ELISA

The conditioned media from the microglia was collected at the end of the experiment and analysed to determine Il1b levels. IL1b levels were assayed in cell media using a commercial ELISA kit (88-7013-88, Thermofisher) following manufacturer’s instructions.

### Statistics

Statistical analysis was performed using Student’s t-test or One-Way ANOVA using Graphpad. Results are expressed as the mean ± S.E.M. A p<0.05 was considered significant as the following: * p<0.05; ** p<0.01; *** p<0.001

